# Full-duplex acoustic interaction system for cognitive experiments with cetaceans

**DOI:** 10.1101/2022.05.27.493738

**Authors:** Jörg Rychen, Julie Semoroz, Alexander Eckerle, Richard HR Hahnloser, Rébecca Kleinberger

## Abstract

Cetaceans show high cognitive abilities and strong social bonds. Acoustics is their primary modality to communicate and sense the environment. Research on their echolocation and vocalizations with conspecifics and with humans typically uses visual and tactile systems adapted from research on primates or birds. Such research would benefit from a purely acoustic communication system in which signals flow in both directions simultaneously. We designed and implemented a full duplex system to acoustically interact with cetaceans in the wild, featuring digital echo-suppression. We pilot tested the system in Arctic Norway and achieved an echo suppression of 18 dB leaving room for technical improvements addressed in the discussion. Nevertheless, the system enabled vocal interaction with the underwater acoustic scene by allowing experimenters to listen while producing sounds. We describe our motivations, then present our pilot deployment and give examples of initial explorative attempts to vocally interact with wild orcas and humpback whales.

## Introduction

Dolphins have long been a subject of interest for researchers due to their outstanding cognitive abilities and communicative skills [1], [2]. Studying dolphins through direct human interactions presents unique challenges, especially with wild animals. The aquatic environment is restrictive of sustained interactions, and the air/water barrier hampers acoustic communication. Early experiments in 1978 by John Lilly [3] aimed to establish human-dolphin communication by co-living a human experimenter and a dolphin in a specially designed house, but the scientific outcome remained limited. A few years later, experiments showed that dolphins can understand the syntax and semantics of visual and acoustic commands [4], suggesting complex auditory learning through reinforcement by food rewards. Vocal production learning and the ability to label objects have also been demonstrated [5]. These observations led to research endeavors using custom-built keyboards and touchscreens for the animals. Such paradigms were often adapted from established techniques used for primates and birds. An extensive review of such systems by Denise Herzing [6] provides various insights into the design of technologies for communicating with Delphinidae. Her analysis advocated for approaches that keep the human in the loop, rather than aiming for subjective cognitive assessment. She also recommends approaches based on sound, rather than other senses, as more natural for the animals. She highlighted the lack of tools adapted for underwater communication, emphasizing the importance of two-way audio.

Recent studies have since made stronger use of audio, for instance by using a hydrophone array within a large visual screen such that the dolphins could “point to” a location on the screen with an acoustic click [7]. Denise Herzing studied habituated dolphins in the wild while they interacted with human swimmers equipped with a device that could play back and detect a set of preprogrammed whistles [8]. These experiments addressed the ability of dolphins to use referential signals for objects or actions. By using a two-way acoustic interaction system, the researchers thus were able to conduct these experiments without reinforcement by food reward but were based solely on the curiosity and playfulness of the animals. However, current acoustic systems for marine mammals, even two-way ones, do not allow for communication in both directions simultaneously, which is referred to as full-duplex. In the following section, we highlight the important difference between full-duplex and half-duplex systems.

### Full-duplex communication systems and the need for echo suppression

Full-duplex signal transmission means that signals can be transmitted in both directions simultaneously. An example of a full-duplex system is the analog telephone, where both parties can simultaneously speak and listen. An example of a half-duplex system is the walkie-talkie, whereby a push-to-talk button enables the sender and disables the receiver of the device.

The difficulty of full-duplex systems is to avoid echoes and feedback that can occur when input and output devices are close to each other. When this happens, the signal broadcast over the speaker is picked up by the microphone and transmitted back to the sender. Echoes can be distracting and may lead to feedback oscillations, producing intense sounds. Therefore, the self-produced signal needs to be separated from the received signal at the microphone (Figure 1).

**Figure 1.**
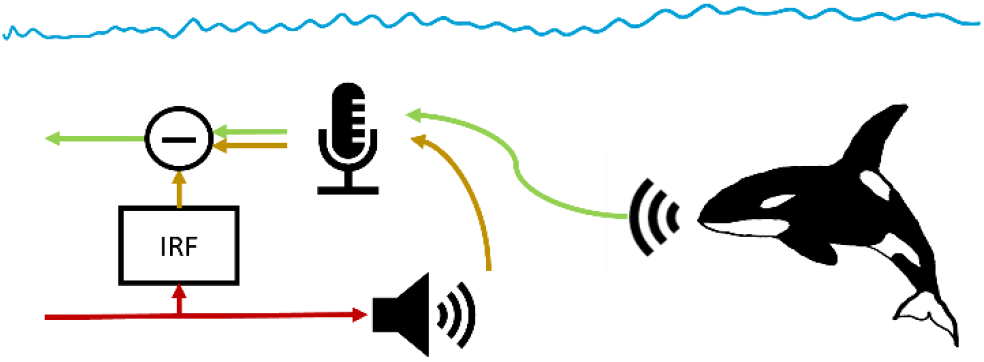
Full-duplex communication system with echo suppression. The signal that is broadcast over the speaker (red) is transferred acoustically to the hydrophone (yellow), where it is superimposed over the external signal of interest (green). Echo suppression is a digital filter that models the transfer of the signal from speaker to microphone with an impulse response function (IRF). This modeled signal is subtracted from the raw microphone signal to yield a pure external signal (green).

Echo suppression (also known as “self-interference suppression”) is a signal-processing technique developed to separate both directions of a signal [9]. It is a key technology for online conference applications and part of mobile phones. Echo suppression requires knowledge of the transfer function from the speaker to the microphone, i.e., described by the impulse response function (IRF), which is the response to a virtual, infinitely sharp impulse in the time domain. Such filters are implemented by adaptive filtering with the least-mean-squares (LMS) algorithm and can be tuned, for example, by playing white noise [10].

Although full-duplex acoustic communication systems with echo suppression are widely used in underwater applications [11], to the best of our knowledge, they have not yet been applied in research on either human-dolphin communication or cetaceans ’cognitive abilities.

We argue that full-duplex communication systems would enhance existing research protocols and enable new research applications. In the following sections, we review four classes of experiments that could benefit from full-duplex audio interactions.

### Interactive Playback Experiments (Figure 2A)

**Figure 2.**
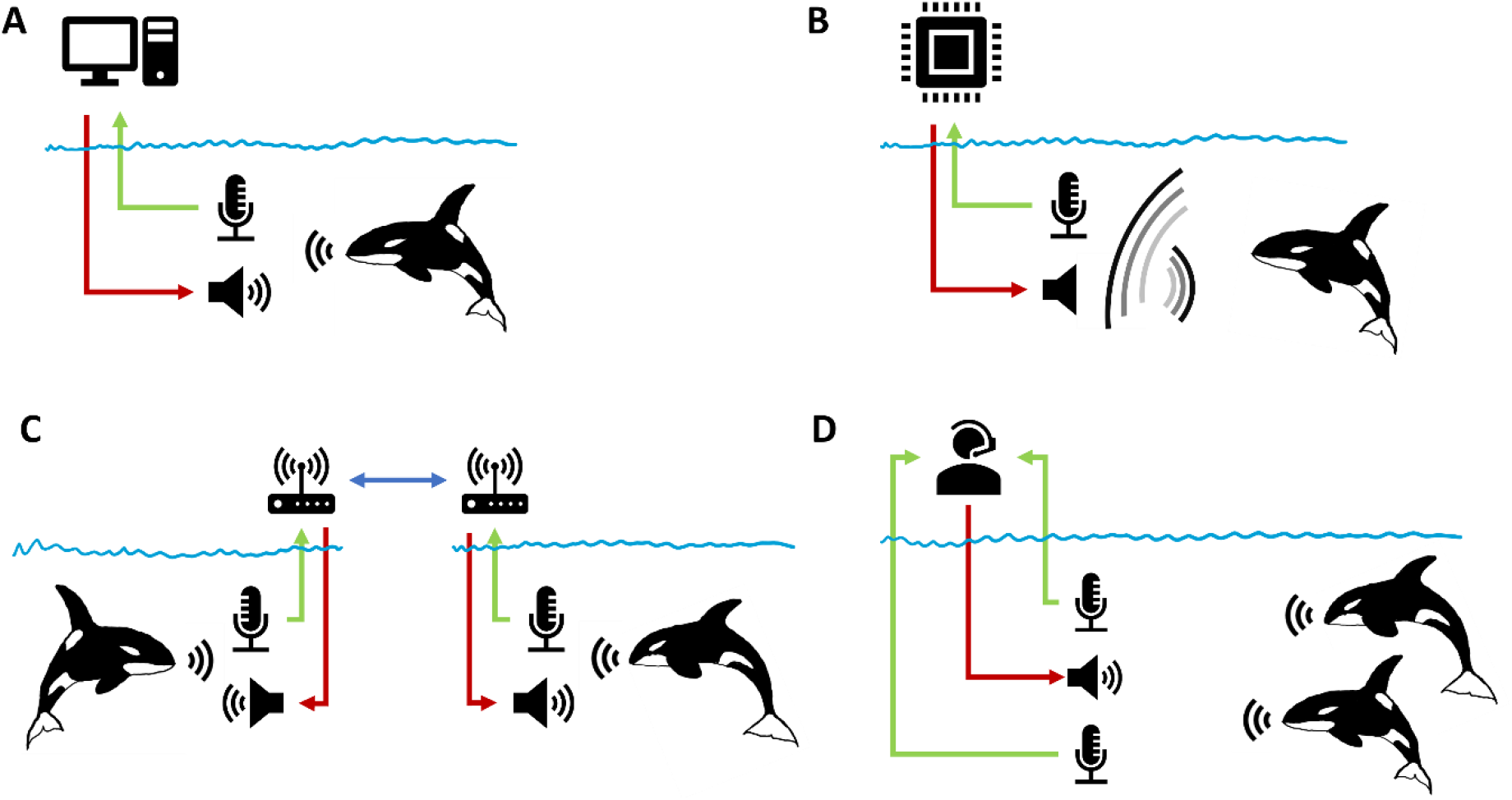
Four classes of acoustic interactive experiments with cetaceans that can benefit from a full-duplex system: **A:** Computerized interactive playback experiments. Prerecorded acoustic stimuli are played back depending on the behavior of the animal. **B**: Phantom echo (artificial sonar target). A low latency digital signal processing unit detects incoming echolocation clicks, and then emits a computed echo to simulate a virtual target. **C:** Telecommunication between two animals to study their communication. This approach provides separated recordings of individuals, the signals can be modified or filtered in real-time, and additional signals can be injected. **D:** Direct human–animal vocal interaction. This setup allows humans to interactively experience the underwater acoustic space.

Playback experiments are an important tool in research of animal communication and serve to validate hypotheses on the function or meaning of acoustic signals [12]. Although these experiments often require ethical considerations to avoid deception, they help to demonstrate the animal’s ability to discriminate between classes of signals [13], [14]. While these experiments have been principally one-way interactive playback experiments, as highlighted in a review article by Stefanie King [15], they promote research on communication and cognitive tasks. In this context, *interactive* generally means that the played-back signal is chosen dependent on the animal’s behavior. For example, the experimenter may observe or listen to the animal and then decide which prerecorded or synthesized stimulus to play back and when to do it. A full-duplex system would be useful in such experimental approaches because they provide better access to turn-taking behavior, to choral behavior, and to simultaneous vocalization.

### Phantom Echo (Figure 2B)

To study the echolocation abilities of cetaceans, researchers have used the technique of phantom echo (or artificial sonar target) [16], [17]. An incoming signal, i.e., a single click of the animal under investigation, is processed with low latency, and a simulated acoustic reflection of the virtual target is played back to the animal. This approach has been used to study how dolphins can discriminate between metallic cylinders whose walls differ in thickness only by a fraction of a millimeter [18]. In their setup, to prevent repeated echoes, the transducer was placed much closer to the dolphin than the receiving hydrophone. The dolphin also had to remain at a precise position defined by a bite plate. In a related experiment, dolphins were able to detect jittering distance variations of an artificial target [19]. An experiment about auditory stream segregation with actively swimming dolphins [20, Ch. VI] used phantom echoes triggered by on-axis clicks toward the target. The system was implemented on a field-programmable gate array (FPGA) with low and precise latencies and used a “push-to-talk” technique, by blanking the received signal during the emission of the phantom echo, to avoid loops of repeated echoes. This limited the repetition rate of clicks for the dolphin. Full-duplex systems, could enable phantom echo experiments with free moving animals and without restrictions on repetition rate, paving the way for more research on the echo location capabilities of dolphins.

### Animal Telecommunication (Figure 2C)

Dolphins use vocalizations and clicks to communicate between conspecifics. In 1965, an experiment recorded the communication between two dolphins kept in separate tanks, connected with an electronic communication system [21]. This system had no echo-suppression, and the receiver and transmitter had to be placed far apart inside each pool to minimize the echo and prevent feedback oscillations. A squelch was also missing and from the background noise the dolphins could infer when the system was turned on. We argue that the technology of echo suppression presented in this paper could lead to vast variations in experiments on the communication of dolphins. For instance, dolphins performing synchronous aerial jumps have been observed to coordinate their behavior with click trains [22]. Could they synchronize their jumps in separate tanks? It has also been shown that two dolphins together could solve cooperative tasks [23], could they collaborate based solely through an acoustic channel? What kind of information would they be able to share with each other to solve increasingly complex problems? Full-duplex systems have provided some answers to such questions in a study of the influence of social learning on auditory discrimination task in songbirds [24]. Also, animal telecommunication systems allow for modifying signals in real time to test hypotheses about the roles of auditory features for communication. The modifications could include adding noise, delays, or filtering out of individual links in communication networks [10]. In the wild, under water telecommunication systems could be a means to assist entrapped cetaceans [25] to find their way back to their conspecifics by establishing an acoustic link between the entrapment bay and the outside group members.

### Human–Animal Interaction (Figure 2D)

In the pilot deployment described below, a direct human-animal interaction was used to assess the performance of our system and to explore our ability to capture the animals’ attention. In contrast to the interactive playback paradigm (Figure 2A), human-animal vocal interaction is technically simpler and more flexible. While a computer program offers a high degree of repeatability, a human in the loop shifts the research focus to softer factors of communication, such as curiosity, bonding, and surprise. In 2008, Rothenberg played clarinet in accompaniment with a singing male humpback whale and suggested interspecies music-making as a potential tool in helping understand the complex communication strategies of cetaceans [26]. Spontaneously mimicking of human vocalization have been observed in cetaceans and reports suggest mutual curiosity to be a driving factor [27]. In addition, having the human reactivity in the loop could help with studying phenomena such as joint rhythm and turn taking [28] that are still difficult to handle with computer programs. A full-duplex communication system is needed to study these phenomena in a shared communication channel.

## Pilot Experiments

We designed, developed, and field-tested a system for acoustic full-duplex communication between a human experimenter and an underwater acoustic space (see Figure 2D). The system was first tested in 2020 and then deployed in 2021 in Norway in waters frequented by killer whales and humpback whales for the initial observation of their reactions and behaviors.

Through a stereo headphone, the experimenter could listen to the underwater sounds captured by two hydrophones in the water. The experimenter’s vocalizations were broadcast over an underwater speaker located approximately six meters below the boat. Two echo-suppression filters reduced the signals of the direct sound transmission from the speaker to the hydrophones to attenuate the experimenter’s own sounds, and provide better perception of faint distant sounds, and reverberations in the fjords.

The goal of this initial pilot deployment was to collect insights regarding the following three objectives:

1. Assess the performance of our echo-suppression system for underwater acoustics by measuring the suppression of the direct echo in decibels (dB) and identify points of improvement.
2. Listen to wild animals’ behaviors and attempt to react live to their vocalization. Observe potential initial signs of curiosity or reaction of the animals to human-produced sounds
3. Subjectively report on human’s active experience of the underwater acoustic scene.

### System

The system comprised an underwater loudspeaker with a fixed array of four hydrophones (Figure 3B). The electronic system was mounted inside a transport case (Figure 3C), including a battery, analog amplifiers, and a digital signal processing system based on FPGAs (Figure 4). To ensure high dynamic range, low noise, and good linearity, we used instrument-grade analog to digital converters (ADC) and digital to analog converters (DAC) designed for audio applications. The vocalizations of the human experimenter were either captured with a directional microphone or a contact microphone in the form of a piezo disk pressed to the throat.

**Figure 3.**
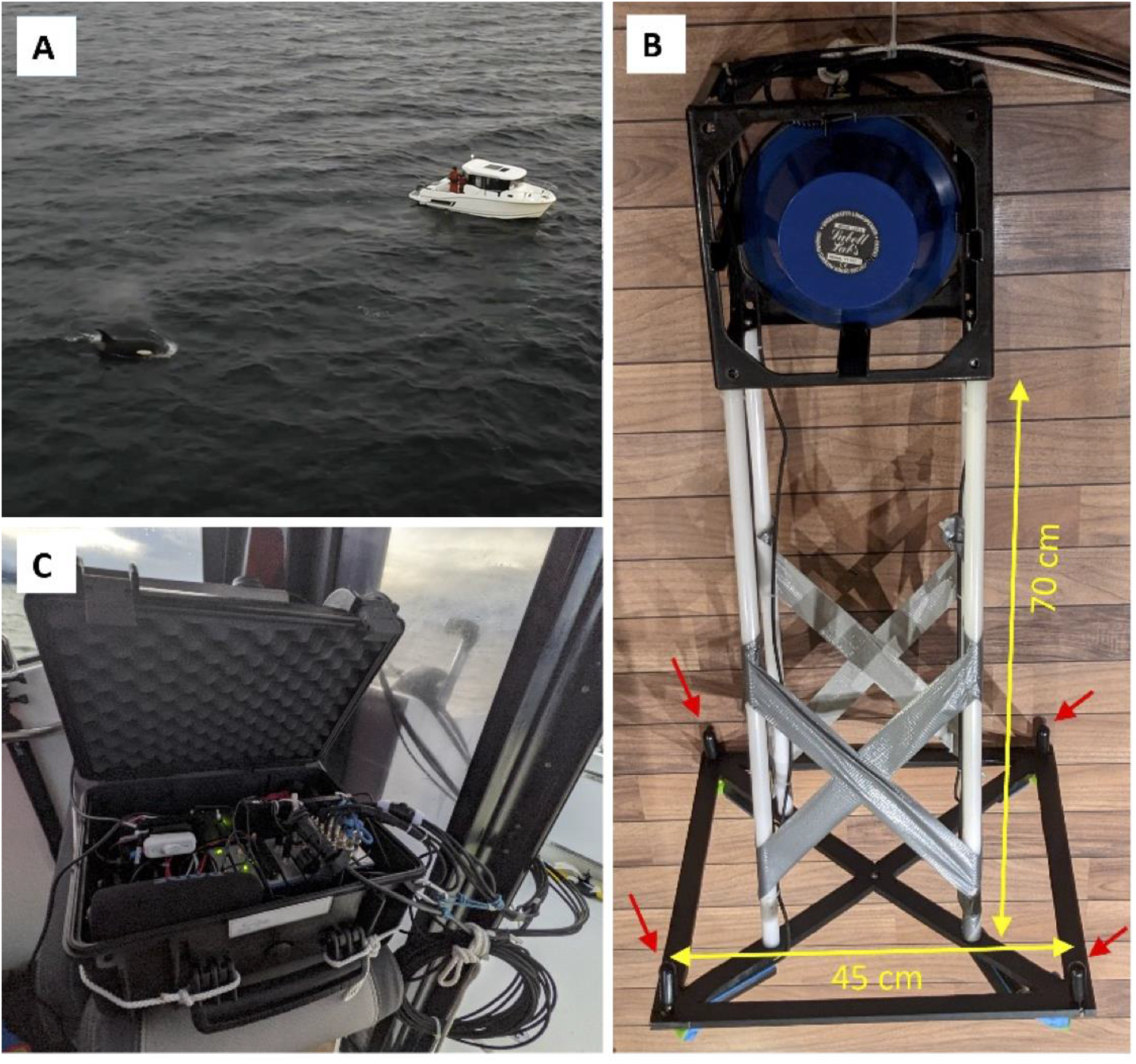
**A**: We installed the full-duplex interaction system on a small motorboat. The experimenter wore a headset and interacted with the underwater sonic environment. **B**: We mounted an underwater loudspeaker (LL916, Lubell Labs, USA) and four hydrophones (HTI-96, High Tech Inc., USA, red arrows) 70 cm below the speaker to a plastic frame made of polyethylene (PE-HD), that minimizes acoustic disturbances). The speaker was lowered overboard to about 6 m into the water. **C**: To protect the electronics from saltwater splashes, they were placed inside the cabin of the motorboat.

**Figure 4:**
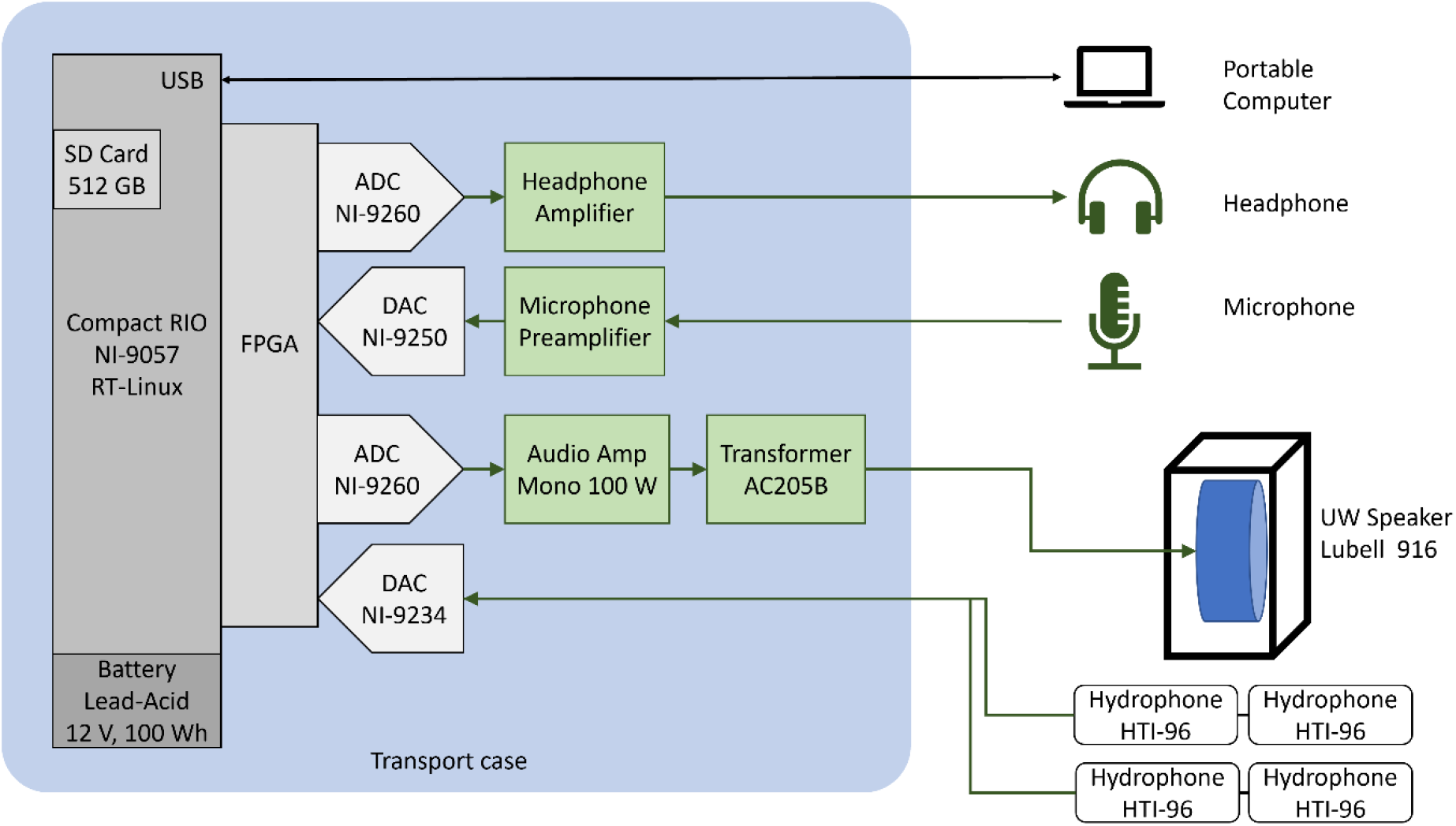
Overview of the full-duplex system architecture. We used a controller (CompactRio, NI-9057, National Instruments, USA) with a real-time Linux operating system. The integrated FPGA interfaces with input and output (IO) modules and runs the signal processing including the adaptive LMS filters for echo-suppression. Digital-to-analog converters (DAC) and analog-to-digital converters (ADC) with 24-bit sigma-delta converters were run at a sample rate of 51.2 kHz. All IO modules were driven by a common clock for synchronized sampling of all channels. The two output channels of the first NI-9260 module were amplified and connected to an analog stereo headphone. A directional condenser microphone was connected to a microphone preamplifier and then to the analog input module NI-9250. The underwater loudspeaker (Lubell Labs, USA) was driven by a mono 100-W class-D amplifier (TPA3116D2, Texas Instruments), followed by an impedance matching circuit (Transformer Model AC202B, Lubell Labs, USA). The signal was generated by one channel of the second NI-9260 module. We used four hydrophones (High Tech Inc., USA) with integrated ICP preamplifiers. These were connected directly to the four input channels of the NI-9234 module, which delivered an excitation current of 2 mA to each of the preamplifiers. We used the factory calibration measurement for hydrophones with sensitivities of −170 dB re 1 V/μPa (+/- 0.3 dB). The system used about 10 W and the 12 V, 100 Wh battery allowed for a run-time of at least 6 h. All signals on the FPGA were saved to a file on an SD-card. Live inspection of the signals and device control was made possible thanks to a user interface running on a portable computer. All programming was done in LabVIEW (National Instruments, USA).

We implemented echo suppression with two adaptive LMS filters of 512 samples length, corresponding to 10.24 ms and to 15.36 m sound propagation distance. The filters were tuned by playing white noise for a few seconds at a source level of 145–155 dB (re μPa rms @ 1 m) in the 200 Hz–20 kHz band. The normalized learning rate was typically 0.01, see [10] for details. We calibrated the loudspeaker and its amplifier based on the frequency-averaged sound-pressure measurement of 184 dB re μPa/V at a distance of 1 m in the axial direction of the speaker in the band of 200 Hz–20 kHz.

### Field Test

We tested the system in December 2021 in fjords near the village of Skjervøy in northern Norway (N 70.0336, E 20.9880), during the season where many humpback whales (*Megaptera novaeangliae*) and killer whales (*Orcinus orca*) regularly feed on herring (Clupea harengus). Using a small motorboat (Figure 3A), we conducted daily excursions looking either for animals or a quiet place. Before recording, we stopped the boat and turned off the engines and the depth finder, lowered our hydroacoustic system into the water, and tuned the echo-suppression filters. Thereafter, wearing a stereo headphone, the experimenter could listen for underwater sounds, including the echoes from self-generated sounds broadcast through the speaker.

### Results and discussion

#### Echo suppression performance

We established an echo suppression of 18 dB, measured with white noise within the 200 Hz to 20 kHz band. The impulse response function is shown in Figure 5A.

**Figure 5.**
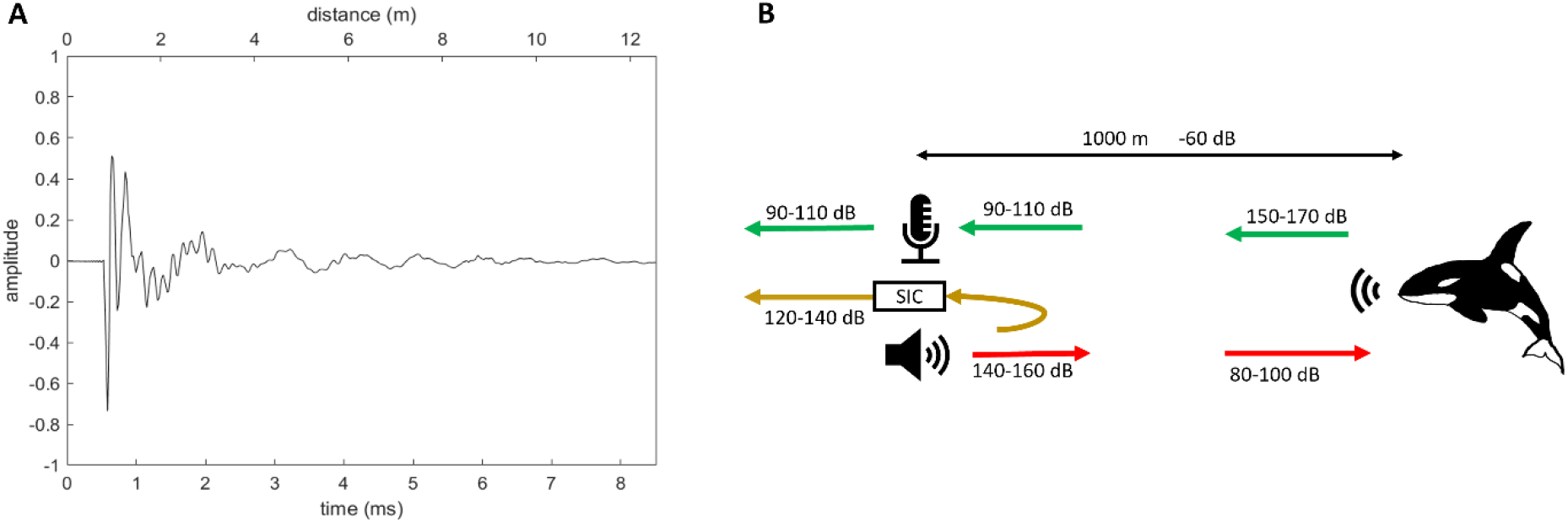
Experimental echo suppression performance. **A**: The impulse response function after tuning the adaptive filter with white noise. The curve represents the 512 filter coefficients of the adaptive filter. The lower x-axis indicates the time delay, and the upper x-axis indicates the corresponding travel distance of the sound waves (c=1475 m/s). **B:** Diagram of sound levels (rms re 1 μPa) in a typical situation with an animal 1 km away. With an exemplary echo-suppression of −20 dB, the own sounds on the headphones are loader than those of the remote animal.

The impulse response function in Figure 5A is zero for the first 0.5 ms, which agrees well with the 70 cm distance between the hydrophones and the speaker (we corrected an offset of 1.388 ms to account for the specified delay of the ADC and DAC and an internal delay of four samples (80 μs) in our signal processing on the FPGA).

Figure 5B shows a hypothetical situation with an animal at a 1km distance, corresponding to an attenuation of −60 dB. Typically, orcas vocalize with a source level of 150-170 dB [29], yielding a sound pressure level of 90-110 dB at the hydrophone. The source level of our speaker was about 140-160 dB, arriving at the animal’s location at a level of 80–100 dB. The echo suppression reduced the level of our own signal to 120-140 dB and thus experimenters typically heard their own voice louder than that of the animals’.

The achieved echo suppression of 18 dB proved less effective than expected. The reason for the limitation in performance is unclear in this case. Typically, nonlinearities of the amplifier and the speaker cause harmonic distortions, which are the main limiting factors of echo suppression systems [30]. A major improvement can be expected by sensing the speaker’s signal with an additional analog-to-digital converter. This signal could be used in the regression of the adaptive filter, circumventing the non-linearity of the power amplifier and transformer. It has been shown that an echo-suppression of 69 dB can be achieved in this way [11]. Another factor limiting the echo suppression is the time-varying impulse response function from a moving sea surface and was addressed in ref [31]. Yet another improvement could be to use a more rigid frame for the loudspeaker and hydrophones (see Figure 3) to avoid mechanical resonances.

This pilot experiment allowed us to assess echo-suppression performance in a realistic experimental situation in a harsh environment.

#### Human experience of underwater soundscape

One motivation behind the development of the echo suppression system was to enable a new form of underwater acoustic perception for humans. The full-duplex system allows for experimenters to be vocalizing and clearly perceiving the echoes and reverberations of their voices within the fiords. The deep and granite-walled fjords of Arctic Norway form an unusual underwater acoustic scene. The echo-suppression filters attenuated echoes from reflections up to 6 m around the speaker (Figure 5A), but further echoes remained unchanged. By using two hydrophones spaced about 45 cm apart, we obtained a stereo percept. The speed of sound being five times faster in water than in air, therefore, from the perspective of interaural time differences, placing the hydrophones 1 m apart in water, corresponds to the roughly 20 cm distance of human ears. To obtain also interaural intensity differences, one would need a heavy and five times larger model of a human head with hydrophones in the ears (e.g., a concrete sculpture). However, computational methods exist to simulate head-related transfer functions, and underwater spatial audio has been demonstrated with a four-hydrophone array [32].

The limited echo-suppression impeded on the perception of distant echoes. The direct signal from the speaker to the hydrophone was strong, limiting the gain at which we could listen to the hydrophones. For a more immersive experience of the of the underwater world, further improvements to the echo-suppression system are needed, which seams feasible [11].

Making the underwater acoustic world perceivable to humans could be an important factor for raising awareness of the unique acoustical world inhabited by these animals, who evolved to use sound as their primary sense. Acoustic pollution is a major thread for killer whales and other cetaceans [33].

#### Vocal interactions with orcas and humpbacks

One important motivation of this research is to enable direct acoustic streaming between cetaceans and human experimenters. During the pilot deployment of the system, we seek for initial interactions with wild animals.

We aimed to engage orcas or humpback whales in interaction by imitating their sounds and adapting to their vocal rhythms. The goal was to address an animal by providing a stimulus that was not random but that had a correlation with its own vocalizations. We expected to observe interest of the animals and playful exploration of the new sound source due to their natural curiosity.

Although, this pilot experiment aimed at testing the new system in realistic conditions and was not designed to gain scientific insights about the animals’ reaction, our initial observations may suggest hints of reaction of the animals to our sounds. Those initial observations are reported bellow.

Figure 6 shows four example spectrograms with animal vocalizations and human sounds in different colors. Subfigures A and C show interactions with humpback whales, and subfigures B and D show interactions with orcas.

**Figure 6:**
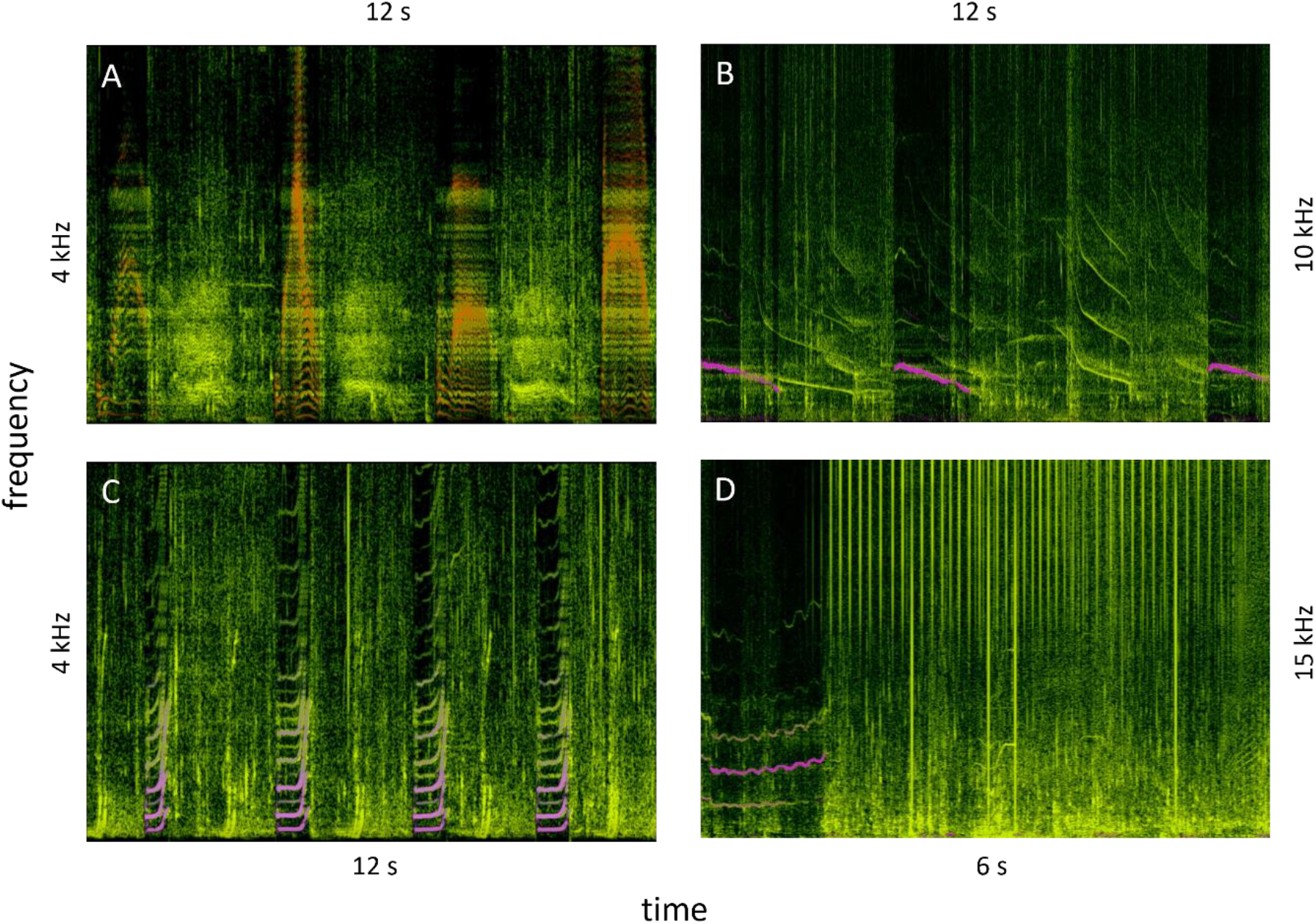
Four example spectrograms of interactions with humpback and killer whales. The horizontal axis is time, with extent indicated in seconds. The vertical axis is linear in frequency starting from zero with extent indicated in kHz. Green-yellow is the signal from the hydrophones and indicate all background sounds including animal vocalization. Overlaid in purple is the human signal broadcast through the speaker. **A**: Humpback growls alternating with human quacks. **B:** Down sweep chorus with orcas. **C**: Rhythmic humpback whups alternating with human whup imitations. **D**: Orca click trains presumably aimed at our acoustic system right after a human high-frequency sound. Some of the clicks overloaded the signal (vertical lines down to the lower edge).

The humpback whales in northern Norway become vocally very active by the end of December. They produce rhythmic sounds, but the sequences are not as stereotyped as those songs on the breeding grounds [34], [35]. They repeat characteristic vocal units, typically 2 – 6 times with gaps of about 2 s between each repeated unit. In some of our encounters, when we tried to produce imitations of their vocalizations during the gaps, they repeated the same unit more than 10 times.

For the orcas, we preferred to experiment with them immediately after a feeding event, when they engage in social and playful behaviors, and disperse into smaller groups, while vocalize extensively. Their whistles are typically up-sweeps, with a few rare instances of reported down-sweeps calls. In a call-type catalog for Norwegian killer whales [36], we found only one call, the N64, with a clear down-sweep frequency contour. In one of our encounters, the experimenter was whistling with down-sweeps and the orcas engaged in a chorus of down-sweep whistles that lasted several minutes. A spectrogram is depicted in Figure 6B. Figure 6D shows an encounter with an immature orca and an adult orca, probably a young-mother pair. The pair approached and circled our boat seemingly investigating our acoustic device in the water with direct click trains, often directly after we broadcast a vocalization. This may suggest that the orcas investigated the new unknown sound source in the water. More investigations based on scientific protocols are needed to validate these observations. Future work includes the identification of measurable factors and protocol designs using the systems focused on call timing, acoustic characteristics, animal location and movements, etc.

## Conclusion

We presented the design and pilot deployment of a full-duplex communication system to research cetaceans’ communication abilities. Our system provides a bidirectional acoustic link into the water. At the heart of the system is the echo-suppression that achieved a performance of 18 dB. We tested the system in northern Norway and made an initial exploratory attempt to vocally interact with wild orcas and humpbacks aiming to engage the animals by imitating their sounds and adapting their rhythm. Further engineering efforts are needed to improve the echo-suppression and adapt the system for specific experimental needs. Such a system may open the door to new research on interactive playbacks, artificial sonar targets, animal telecommunication, and human-animal communication.

## Ethical Statement

We received confirmation from the Norwegian Food Safety Authority that under Norwegian and European legislation related to animal research, formal approval and a license are not required (regulation of 18 June 2015 No 761 concerning the use of animals for scientific purposes § 2, f). The experiment was regarded as non-invasive. Nevertheless, we took technical measures to prevent startling the animals by limiting the rate of onset of acoustic emission levels. We limited the emission level to 170 dB re 1μPa rms. We limited our sonic interactions to periods of 15 minutes.

## Acknowledgments

We would like to thank Fiona Wüthrich, Elizabeth Ren, Berenice Fischer and Linus Rüttimann for their help during the expedition.

## Author contributions

Conceptualization: JR, RK, RH, JS. Methodology: JR, RK. Investigation: JR, JS, AE. Data curation: JS, JR. Software & Hardware: JR, RH. Visualization: JR. Writing – original draft: JR, RK. Writing – review & editing: JR, RK, RH, AE, JS.

## References

[1] D. L. Herzing and C. M. Johnson, Dolphin communication and cognition : past, present, and future / edited by Denise L. Herzing and Christine M. Johnson. The MIT Press, 2015.

[2] L. Marino, “Convergence of complex cognitive abilities in cetaceans and primates,” in Brain, Behavior and Evolution, 2002, vol. 59, no. 1–2, pp. 21–32. doi: 10.1159/000063731.

[3] J. C. Lilly, “Communication between man and dolphin: The possibilities of talking with other species.,” Crown Publishers, Inc., New York. *269pp. 1978*. Crown, New York, p. 269, 1978.

[4] L. M. Herman, D. G. Richards, and J. P. Wolz, “Comprehension of sentences by bottlenosed dolphins,” Cognition, vol. 16, no. 2, pp. 129–219, 1984, doi: 10.1016/0010-0277(84)90003-9.

[5] D. G. Richards, J. P. Wolz, and L. M. Herman, “Vocal mimicry of computer-generated sounds and vocal labeling of objects by a bottlenosed dolphin, Tursiops truncatus.,” J Comp Psychol, vol. 98, no. 1, pp. 10–28, 1984, doi: 10.1037/0735-7036.98.1.10.

[6] D. Herzing, “Interfaces and Keyboards For Human-Dolphin Communication: What Have We Learned?,” Animal Behavior and Cognition, vol. 3, no. 4, pp. 243–254, 2016, doi: 10.12966/abc.04.11.2016.

[7] M. Amundin, J. Starkhammar, M. Evander, M. Almqvist, K. Lindström, and H. W. Persson, “An echolocation visualization and interface system for dolphin research,” J Acoust Soc Am, vol. 123, no. 2, pp. 1188–1194, 2008, doi: 10.1121/1.2828213.

[8] D. Kohlsdorf, S. Gilliland, P. Presti, T. Starner, and D. Herzing, “An underwater wearable computer for two way human-dolphin communication experimentation,” in ISWC 2013 - Proceedings of the 2013 ACM International Symposium on Wearable Computers, 2013, pp. 147–148. doi: 10.1145/2493988.2494346.

[9] M. M. Sondhi, “An Adaptive Echo Canceller,” Bell System Technical Journal, vol. 46, no. 3, pp. 497–511, 1967, doi: 10.1002/j.1538-7305.1967.tb04231.x.

[10] J. Rychen, D. I. Rodrigues, T. Tomka, L. Rüttimann, H. Yamahachi, and R. H. R. Hahnloser, “A system for controlling vocal communication networks,” Scientific Reports, vol. 11, no. 1, pp. 1–15, 2021, doi: 10.1038/s41598-021-90549-0.

[11] L. Shen, B. Henson, Y. Zakharov, and P. Mitchell, “Digital Self-Interference Cancellation for Full-Duplex Underwater Acoustic Systems,” IEEE Transactions on Circuits and Systems II: Express Briefs, vol. 67, no. 1, pp. 192–196, 2020, doi: 10.1109/TCSII.2019.2904391.

[12] V. B. Deecke, “Studying Marine Mammal Cognition in the Wild: A Review of Four Decades of Playback Experiments,” Aquatic Mammals, vol. 32, no. 4, pp. 461–482, 2007, doi: 10.1578/am.32.4.2006.461.

[13] O. A. Filatova, I. D. Fedutin, A. M. Burdin, and E. Hoyt, “Responses of Kamchatkan fish-eating killer whales to playbacks of conspecific calls,” Mar Mamm Sci, vol. 27, no. 2, pp. 26–42, 2011, doi: 10.1111/j.1748-7692.2010.00433.x.

[14] C. Curé et al., “Evidence for discrimination between feeding sounds of familiar fish and unfamiliar mammal-eating killer whale ecotypes by long-finned pilot whales,” Animal Cognition, vol. 22, no. 5, pp. 863–882, 2019, doi: 10.1007/s10071-019-01282-1.

[15] S. L. King, “You talkin’ to me? Interactive playback is a powerful yet underused tool in animal communication research,” Biology Letters, vol. 11, no. 7, 2015, doi: 10.1098/rsbl.2015.0403.

[16] R. Aubauer and W. W. L. Au, “Phantom echo generation: A new technique for investigating dolphin echolocation,” J Acoust Soc Am, vol. 104, no. 3, pp. 1165–1170, 1998, doi: 10.1121/1.424324.

[17] M. W. Muller, W. W. L. Au, P. E. Nachtigall, J. S. Allen, and M. Breese, “Phantom echo highlight amplitude and temporal difference resolutions of an echolocating dolphin, Tursiops truncatus,” J Acoust Soc Am, vol. 122, no. 4, pp. 2255–2262, 2007, doi: 10.1121/1.2769973.

[18] B. K. Branstetter et al., “Spectral cues and temporal integration during cylinder echo discrimination by bottlenose dolphins (Tursiops truncatus),” J Acoust Soc Am, vol. 148, no. 2, pp. 614–626, 2020, doi: 10.1121/10.0001626.

[19] J. J. Finneran et al., “Dolphin echo-delay resolution measured with a jittered-echo paradigm,” J Acoust Soc Am, vol. 148, no. 1, pp. 374–388, 2020, doi: 10.1121/10.0001604.

[20] C. Malinka, “Biosonar of narrow-band high-frequency toothed whales: Sampling a dynamic, multi-target world,” Aarhus University, 2021.

[21] T. G. Lang and H. A. P. Smith, “Communication between dolphins in separate tanks by way of an electronic acoustic link,” Science Science, vol. 150, no. 3705, pp. 1839–1844, 1965, doi: 10.1126/science.150.3705.1839.

[22] J. Lopez Marulanda et al., “Acoustic behaviour of bottlenose dolphins under human care while performing synchronous aerial jumps,” Behavioural Processes, vol. 185, p. 104357, Apr. 2021, doi: 10.1016/j.beproc.2021.104357.

[23] S. L. King, E. Guarino, K. Donegan, C. McMullen, and K. Jaakkola, “Evidence that bottlenose dolphins can communicate with vocal signals to solve a cooperative task,” Royal Society Open Science, vol. 8, no. 3, 2021, doi: 10.1098/rsos.202073.

[24] G. Narula, J. A. Herbst, J. Rychen, and R. H. R. Hahnloser, “Learning auditory discriminations from observation is efficient but less robust than learning from experience,” Nature Communications, vol. 9, no. 1, p. 3218, 2018, doi: 10.1038/s41467-018-05422-y.

[25] E. Jourdain et al., “Natural Entrapments of Killer Whales (Orcinus orca): A Review of Cases and Assessment of Intervention Techniques,” Frontiers in Conservation Science, vol. 2, no. August, pp. 1–13, 2021, doi: 10.3389/fcosc.2021.707616.

[26] D. Rothenberg, “Whale music: Anatomy of an interspecies duet,” Leonardo Music Journal, vol. 18, pp. 47–53, 2008, doi: 10.1162/lmj.2008.18.47.

[27] S. Ridgway, D. Carder, M. Jeffries, and M. Todd, “Spontaneous human speech mimicry by a cetacean,” Current Biology, vol. 22, no. 20. Elsevier, pp. R860–R861, 2012. doi: 10.1016/j.cub.2012.08.044.

[28] S. Pika, R. Wilkinson, K. H. Kendrick, and S. C. Vernes, “Taking turns: Bridging the gap between human and animal communication,” Proceedings of the Royal Society B: Biological Sciences, vol. 285, no. 1880, 2018, doi: 10.1098/rspb.2018.0598.

[29] M. M. Holt, D. P. Noren, and C. K. Emmons, “Effects of noise levels and call types on the source levels of killer whale calls,” J Acoust Soc Am, vol. 130, no. 5, pp. 3100–3106, 2011, doi: 10.1121/1.3641446.

[30] F. Ling, “Achievable performance and limiting factors of echo cancellation in wireless communications,” 2014. doi: 10.1109/ITA.2014.6804210.

[31] L. Shen, Y. Zakharov, B. Henson, N. Morozs, and P. D. Mitchell, “Adaptive filtering for full-duplex UWA systems with time-varying self-interference channel,” IEEE Access, vol. 8, pp. 187590–187604, 2020, doi: 10.1109/ACCESS.2020.3031010.

[32] S. Delikaris-Manias, L. McCormack, I. Huhtakallio, and V. Pulkki, “Real-time underwater spatial audio: a feasibility study,” 144th Audio Engineering Society Convention 2018, no. May, 2018.

[33] E. Jourdain et al., “North Atlantic killer whale Orcinus orca populations: a review of current knowledge and threats to conservation,” Mammal Review, vol. 49, no. 4, pp. 384–400, 2019, doi: 10.1111/mam.12168.

[34] S. Handel, S. K. Todd, and A. M. Zoidis, “Hierarchical and rhythmic organization in the songs of humpback whales (Megaptera novaeangliae),” Bioacoustics, vol. 21, no. 2, pp. 141–156, 2012, doi: 10.1080/09524622.2012.668324.

[35] E. E. Magnúsdóttir and R. Lim, “Subarctic singers: Humpback whale (Megaptera novaeangliae) song structure and progression from an Icelandic feeding ground during winter,” PLoS ONE, vol. 14, no. 1, 2019, doi: 10.1371/journal.pone.0210057.

[36] A. D. Shapiro, “Orchestration : the movement and vocal behavior of free-ranging Norwegian killer whales (Orcinus orca),” MASSACHUSETTS INSTITUTE OF TECHNOLOGY, 2008. doi: 10.1575/1912/2421.

